# pH-dependent regulation in SLC38A9

**DOI:** 10.1101/2025.10.24.684471

**Authors:** Xuelang Mu, Ampon Sae Her, Tamir Gonen

## Abstract

Cells rely on precise metabolic control to adapt to environmental cues. The mechanistic target of rapamycin complex 1 (mTORC1) senses nutrient availability, with amino acids serving as key signals. Lysosomes, which act as nutrient recycling centers, maintain amino acid homeostasis by breaking down macromolecules and releasing amino acids for cellular use. SLC38A9, a lysosomal amino acid transporter, functions as both a transporter and a sensor in the mTORC1 pathway. Here, we investigated whether SLC38A9 activity is regulated by pH. We show that arginine uptake by SLC38A9 is pH-dependent, and that the histidine residue His544 serves as the pH sensor. Mutating His544 abolishes the pH dependence of arginine uptake without impairing overall transport activity, indicating that His544 is not directly involved in substrate binding. Instead, protonation or deprotonation of His544 appears to influence transport through SLC38A9. To explore this mechanism, we compared two structures of SLC38A9 that we determined, one at high pH and one at low pH, and proposed a working model for pH-induced activation. These findings highlight the role of local ionic changes in modulating lysosomal transporters and underscore the intricate regulatory mechanisms that govern SLC38A9 function and, ultimately, mTORC1 signaling.

## Introduction

Sensing the environment is essential for cells to coordinate their metabolic activities. Cells must respond to diverse cues, including nutrients, energy levels, and growth factors. A central regulator of this process is the mechanistic target of rapamycin complex 1 (mTORC1), a key hub in cellular metabolism. mTORC1 integrates fluctuations in extracellular and intracellular nutrients and directs cellular outcomes, promoting anabolic processes such as protein, lipid, and nucleotide synthesis, or initiating catabolic processes such as lysosome biogenesis and autophagy[1]. Among these signals, amino acids serve as fundamental cues for the mTORC1 pathway, and maintaining a balance between amino acid synthesis and degradation is critical[2]. Lysosomes function as recycling centers, generating nutrients and storing amino acids. They sustain a low pH to enable the breakdown of proteins, polysaccharides, lipids, and nucleic acids. Consequently, there is a close relationship between lysosomal pH and its functions. Importantly, many transporters are embedded in the lysosomal membrane and operate within the pH gradient between the lysosome and the cytosol[3–5].

SLC38A9 is a low-affinity lysosomal amino acid transporter that functions as an upstream amino acid sensor for mTORC1 activation[6–8]. In addition to sensing, it mediates the efflux of several essential amino acids, including leucine and glutamine, in an arginine-dependent manner[9]. Thus, SLC38A9 acts as a transceptor, combining the roles of both transporter and receptor[10,11]. Belonging to the amino acid/auxin permease family, it is characterized by 10-11 transmembrane domains[12] and is also classified within the sodium-coupled amino acid transporter family[6,7]. The zebrafish ortholog (Danio rerio; drSLC38A9) shares 61.9% sequence identity and 86.6% similarity with the human protein. Crystal structures of drSLC38A9 revealed 11 transmembrane helices, with the N-terminus facing the cytosol and the C-terminus positioned in the lysosomal lumen[13] The arginine-bound structure identified a substrate-binding pocket buried among the transmembrane helices[13]. In contrast, the arginine-free structure showed the N-terminus forming a β-hairpin that lodged within the transporter and occupied the arginine-binding site[14]. These findings led to a proposed “ball-and-chain” model to explain amino acid sensing and signaling by SLC38A9. Complementary cryo-EM studies of an N-terminal peptide of SLC38A9 bound to Rag GTPases revealed how the N-terminus regulates nucleotide state switching in the GTPases[15]. Together, these results suggest that under nutrient-rich conditions, when luminal arginine levels are high, arginine displaces the N-terminal domain from the binding pocket, freeing it to interact with Rag GTPases and activate mTORC1[14].

Several lysosomal membrane proteins are known to be regulated by pH[16–20]. This regulation is particularly common at the lysosomal membrane due to the steep pH gradient across it: approximately pH 7.2 in the cytoplasm versus pH 4.7 inside the lysosomal lumen[21]. Because SLC38A9 functions at the lysosomal membrane, we hypothesized that it too may be subject to pH regulation. Proteins regulated by pH, such as transporters and pH-sensitive enzymes, often contain amino acid residues with side chains that can donate or accept protons depending on the local microenvironment[22,23]. Among these, histidine (pKa ∼6.2 in solution) is particularly well known for its role in pH sensing[24,25]. The imidazole side chain of histidine can either gain or lose a proton depending on the surrounding pH, enabling histidine residues to act as molecular pH sensors in a wide range of biochemical processes. Indeed, many pH-sensitive systems rely on histidine residues as key regulators[26–31].

In this study, we examined the pH dependence of SLC38A9 transport activity using purified protein reconstituted into vesicles and radiolabeled ligand uptake assays. By measuring [^3^H]-arginine uptake across different pH conditions, we found that SLC38A9-mediated transport is strongly pH-dependent. Sequence analysis combined with site-directed mutagenesis identified His544 as a key pH-sensing residue in human SLC38A9. Based on these results and supporting structural analysis, we propose a working mechanism in which His544 acts as a pH-regulated allosteric site, modulating arginine transport in response to intracellular pH fluctuations.

## Results

### 1. Purified SLC38A9 transports [^3^H]-arginine *in vitro* and is pH regulated

SLC38A9 was expressed and purified as described in the methods section. Purified SLC38A9 was reconstituted into proteoliposomes at a neutral internal buffer (pH_in_ 7.5) and contain an internal sodium concentration of 10mM and potassium concentration of 90mM. This buffer condition mimics the cytosolic environment[21]. Proteoliposomes were then exchanged into an external buffer with acidic pH (pH_out_ 5.5) and a sodium concentration of 100mM, containing [^3^H]-arginine as substrates (Figure 1A). This sets up a pH and salt to facilitate arginine transport via SLC38A9 until saturation was observed at t= 10 minutes (Figure 1B). To investigate the effect of pH on the transport activity of SLC38A9, we measured the uptake of 3H-arginine under different external buffer pH (pH_out_) at pH 5.5, 6.0, 6.5, 7.0, 7.5 and 8.2, while maintaining the inside buffer pH (pH_in_) consistent at pH 7.5. These experiments demonstrated that the uptake of [^3^H]-arginine was highest at the external pH at 5.5 and 6.0 suggesting that SLC38A9 is indeed pH regulated (Figure 1C).

**Figure 1.**
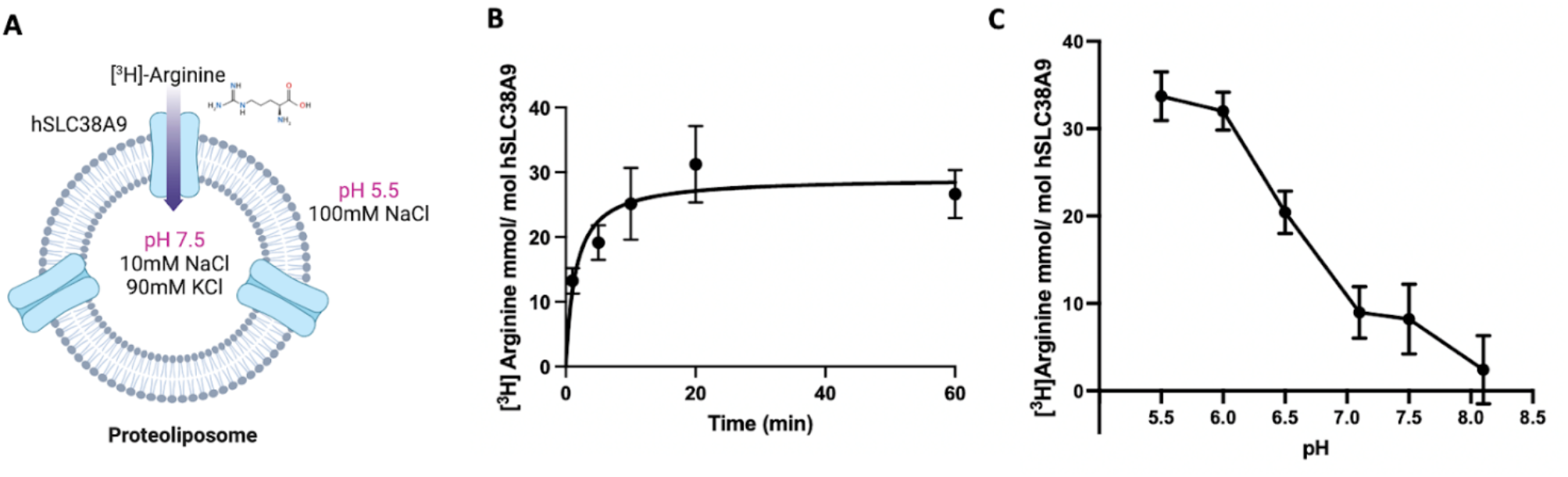
[^3^H]-arginine uptake assays of SLC38A9 proteoliposomes. **(A)** Model representation of SLC38A9 proteoliposomes used in this study. **(B)** Uptake assay in SLC38A9-reconstituted liposomes showing the efficiency of arginine transport with pH_in_= 7.5 and pH_out_= 5.5. This mimics the lysosome-cytosol pH gradients in the cell. Transport was calculated by subtracting the radioactivity associated with empty liposome controls. Error bars, s.e.m. from three independent proteoliposome preparations; *n* = 3 biological replicates. **(C)**Effect of pH on the reconstituted SLC38A9. Reconstitution and transport assay were performed at the indicated pH_out_. Transport was calculated by subtracting the radioactivity associated with empty liposomes. Error bars, s.e.m. from three independent proteoliposome preparations; *n* = 3 biological replicates.

### 2. Histidine 544 is identified as the pH sensor

The identification of a pH sensor is critical for elucidating the molecular mechanism underlying the pH-dependent transport activity of SLC38A9. A *bona fide* pH sensor must undergo pH-induced changes, either to mediate new interactions or to abolish existing interactions, thereby modulating substrate transport. Such pH-induced changes are typically mediated by the protonation and deprotonation of ionizable amino acid residues, such as histidine, glutamic acid, and aspartic acid[23]. Given that SLC38A9 is activated at pH values of 5.5-6.0, the amino acid histidine, with a pK_a_of ∼6.2, emerges as a plausible pH sensor, as it can undergo reversible protonation under these conditions.

We proposed that a functionally important histidine residue may be evolutionarily conserved. We conducted a sequence alignment of SLC38A9 protein sequences from six different species ((*<u>Danio rerio</u>* <u>(Zebrafish)</u>, *<u>Rattus norvegicus</u>* <u>(Rat)</u>,*<u>Mus musculus</u>* <u>(Mouse)</u>, *<u>Homo sapiens</u>* <u>(Human)</u>, *<u>Xenopus laevis</u>* <u>(African clawed frog)</u>, and *<u>Xenopus tropicalis</u>* <u>(Western clawed frog)</u>) to identify conserved histidine residues (Figure 2). Four histidine residues (His339, His373, His480, and His544) within the human SLC38A9 protein are conserved across five different species, as shown in green in the protein sequence alignment (Figure 2). Consequently, these four histidine residues are prime candidates for pH-regulatory functions.

**Figure 2.**
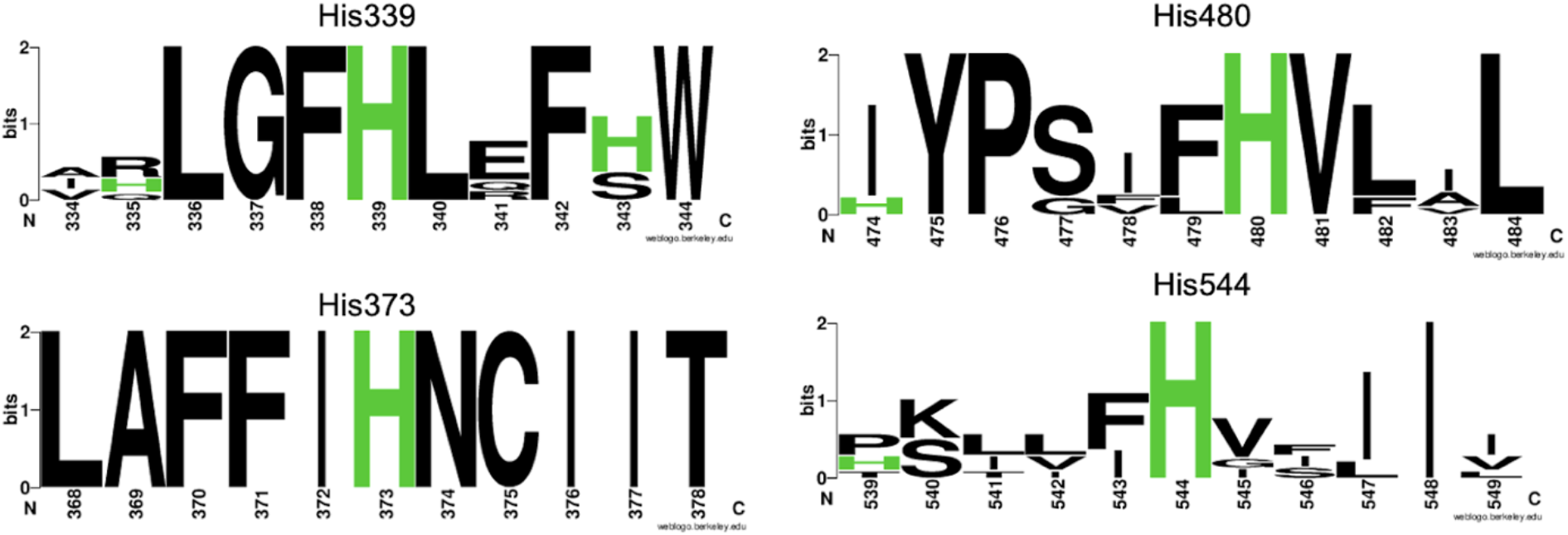
The consensus logo plot for conserved histidine residues. The logo plot was generated from the multiple sequence alignment of 6 species of SLC38A9 from the UniProt database. His339, His373, His480, and His544 are shown to be highly conserved among different species of SLC38A9.

Next, we generated mutants of SLC38A9, in which each of these histidine residues (His339, His373, His480, and His544) were individually mutated to alanine, resulting in mutants designated as H339A, H373A, H480A, and H544A. Importantly, each of these mutants demonstrated functional proficiency, as evidenced by their ability to facilitate the transport of [^3^H]-arginine into proteoliposomes. Importantly, all mutants also displayed pH sensitivity with the exception of H544A. The mutants H339A, H373A, and H480A displayed pH-dependent regulation of [^3^H]-arginine uptake where the maximum activity occurred at the external pH of 5.5. In contrast, when Histidine 544 was mutated to alanine (H544A), the [^3^H]-arginine uptake remained unaffected by external pH variations, indicating the loss of pH-regulatory capabilities in the H544A SLC38A9 mutant (Figure 3A).

**Figure 3.**
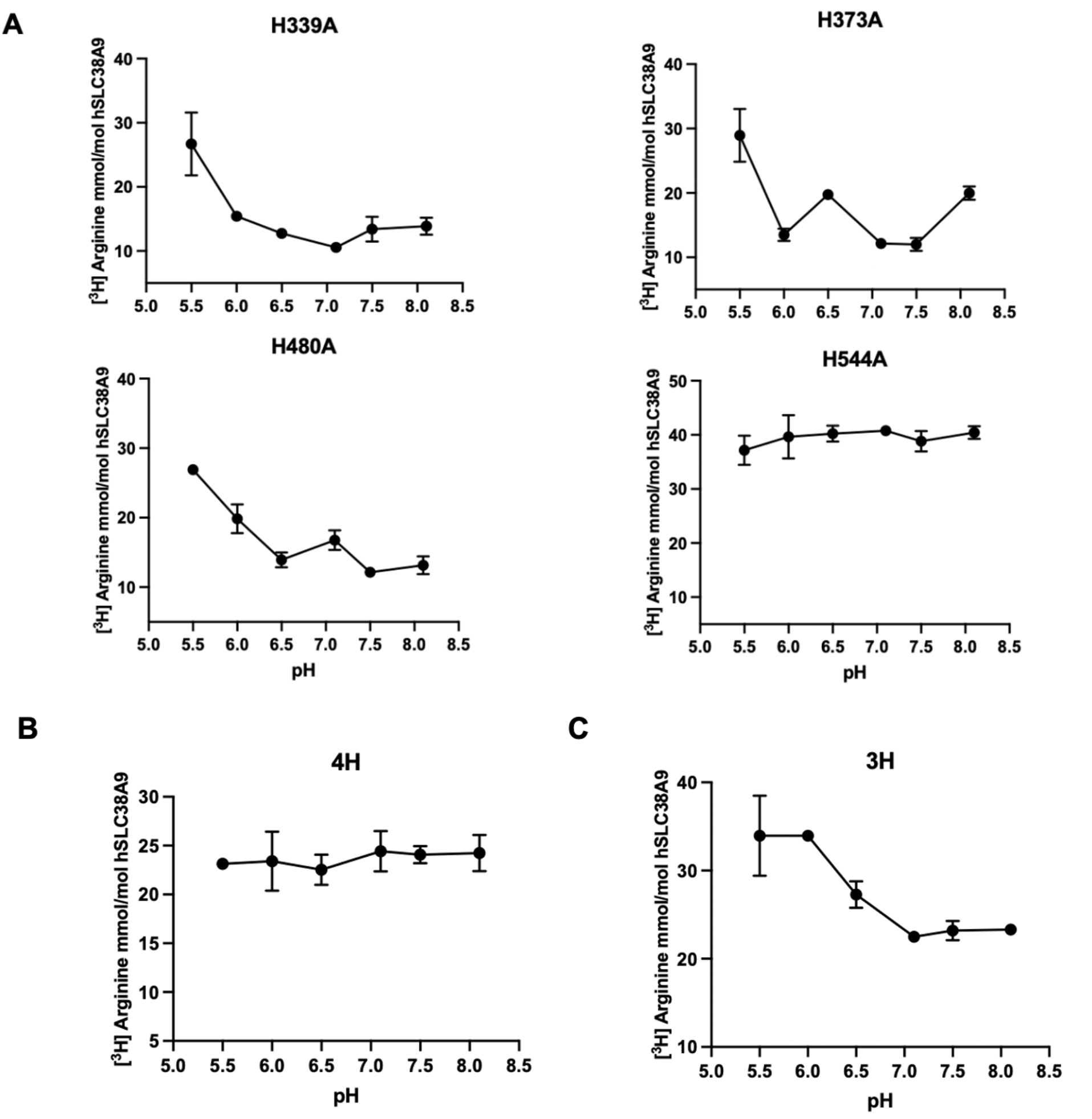
[^3^H]-arginine uptake assays of histidine mutants of SLC38A9. **(A)** Effect of pH on single site mutants of SLC38A9 (H339A, H373A, H480A, and H544A) in reconstituted proteoliposomes. All mutants (H339A, H373S, H480A) but H544A show pH dependency where the maximum activity is at pH 5.5 and decreases with higher pH. Error bars, s.e.m. from three independent proteoliposome preparations; n = 3 biological replicates. Empty liposome controls were subtracted from these final arginine uptake graphs. **(B)** Effect of pH on the quadruple site mutants of SLC38A9 (H339A, H373A, H480A, and H544A) in reconstituted proteoliposomes. **(C)** Effect of pH on the triple site mutants of SLC38A9 (H339A, H373A, and H480A) in reconstituted proteoliposomes. **(D)**Error bars, s.e.m. from three independent proteoliposome preparations; *n* = 3 biological replicates. Empty liposome controls were subtracted from these final arginine uptake graphs.

To further verify whether H544 is the pH sensor, we engineered two additional mutants: a quadruple-site mutant containing all four histidine mutations (i.e. H339A, H373A, H480A, and H544A; referred to as the 4H mutant) and a triple-site mutant comprising H339A, H373A, and H480A mutations (referred to as the 3H mutant). While the 4H mutant showed abolished pH sensitivity the 3H mutant maintained pH sensitivity (Figure 3B and 3C, respectively). These results further suggest that H544 acts as the pH sensor in SLC38A9.

### 3. SLC38A9 is regulated by pH, and coupled to Na^+^ transport

The transport activity of wild type SLC38A9 varied significantly under different pH conditions across liposomal membranes. The uptake of [^3^H]-arginine increased substantially when a pH gradient was present, whereas uptake was minimal in the absence of a pH difference (Figure 4A). To determine whether SLC38A9 is directly coupled to the proton gradient or if its function is merely influenced by pH variations, we examined [^3^H]-arginine uptake in the presence of reversed proton gradients across liposomal membranes. Under an inward proton gradient (pH 7.5 internal/ pH 5.5 external), SLC38A9-mediated uptake of [^3^H]-arginine was observed. Reversing the proton gradient to outward (pH 5.5 internal/ pH 7.5 external) did not abolish [^3^H]-arginine uptake. However, when both internal and external pH were symmetrically set to 7.5 (meaning no pH gradient), [^3^H]-arginine uptake was markedly reduced (Figure 4B). These findings suggest that arginine uptake by SLC38A9 is not dependent on the direction of the proton gradient, as both inward and outward gradients can enhance [^3^H]-arginine uptake.

**Figure 4.**
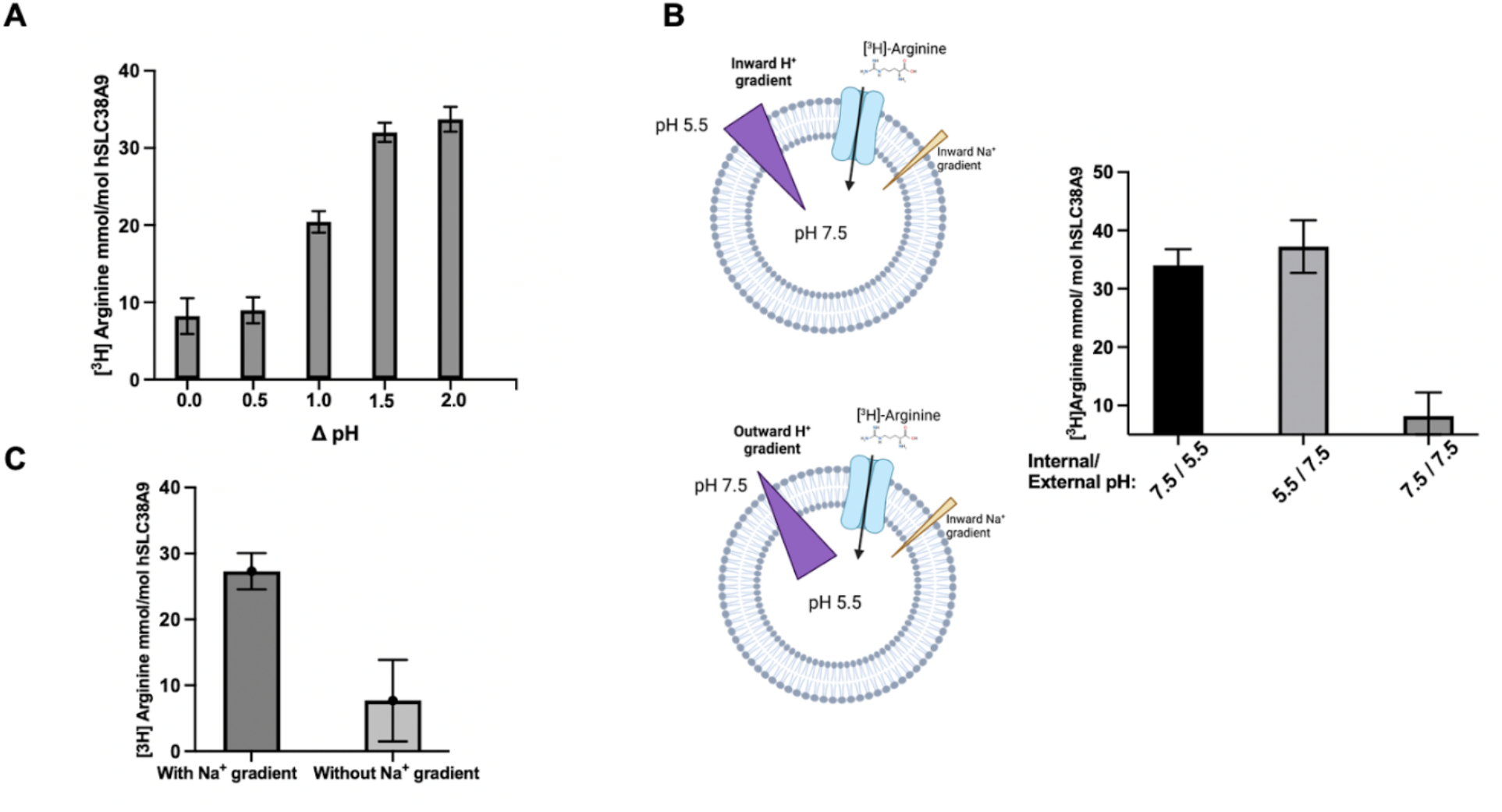
[^3^H]-arginine uptake assays of SLC38A9 proteoliposomes in different pH gradients and Na^+^ gradients. **(A)**The equilibrium uptake of [^3^H]-arginine by SLC38A9 proteoliposomes varied with the differences between pH_in_ and pH_out_. **(B)** [^3^H]-arginine uptake activity of SLC38A9 proteoliposomes in the presence of an inward vs outward H^+^ gradient vs symmetric pH. All the experiments had an inward Na^+^ gradient. **(C)** [^3^H]-arginine uptake assays of SLC38A9 with and without Na^+^ gradient. **(D)** Error bars, s.e.m. from three independent proteoliposome preparations; *n* = 3 biological replicates. Empty liposome controls were subtracted from these final arginine uptake graphs.

Additionally, these results highlight the essential role of an acidic environment on one side of the vacuolar membrane in supporting human SLC38A9 activity. In drSLC38A9, we observed a transport pattern similar to that of SLC38A9. The maximum uptake of [^3^H]-arginine occurred at an internal pH of 7.5 and an external pH of 6.5 (Supplemental Figure 1). However, when the external pH was raised to alkaline levels (at external pH of 7.5 and 8.0), [^3^H]-arginine uptake remained consistently low, regardless of the presence of a proton gradient. These findings confirm that [^3^H]-arginine transport by SLC38A9 is not solely driven by a proton gradient but requires an acidic pH on one side of the membrane for efficient uptake.

SLC38A9 is as a sodium-coupled transporter[6,7]. Previous studies using liposomes reconstituted with drSLC38A9 in various cationic buffers, the presence of sodium significantly enhanced arginine uptake compared to potassium[13]. This observation prompted us to investigate whether SLC38A9 relies on sodium gradients across liposomal membranes for its function. To address this, we established conditions with a low internal sodium concentration (10 mM) and a high external concentration (100 mM). Under these conditions, SLC38A9-reconstituted proteoliposomes exhibited substantial [^3^H]-arginine uptake (Figure 4C). In contrast, when there was no sodium gradient across the liposomes (with 100mM NaCl inside and outside), [^3^H]-arginine uptake was negligible in SLC38A9-containing proteoliposomes, even in the presence of an optimal pH gradient (pH 7.5 inside and pH 5.5 outside) (Figure 4C). These findings indicate that sodium gradients serve as the primary driving force for arginine transport in SLC38A9, while proton gradients function as regulators of transport activity rather than as the driving force.

### 4. pH-sensing mechanism of His544

Our experimental evidence suggests that His544 may function as a pH sensor in SLC38A9 and that protonation or deprotonation of His544 may influence the ability of substrate to bind to SLC38A9 and to be transported.

This hypothesis is supported by structural analysis of drSLC38A9 crystal structure[13,14], one of which was determined at pH 7.2 and the other at pH 6.0. The substrate-bound crystal structure of drSLC38A9, solved at pH 7.2[13], reveals that His532–the homologous residue of human SLC38A9 His544—interacts with Arg155 from TM2 through a cation-pi interaction between the neutral His and the cation Arg^+^ (Figure 5A). In contrast, the drSLC38A9 crystal structure which was obtained at pH 6.0 (an acidic condition that facilitates transport), captured the transporter in a substrate-free conformation[14] without bound arginine. In this state, protonated His532 formed interactions with Tyr151 from TM2 and Ser511 from TM10, indicating a shift in its interaction network compared to the high-pH structure (Figure 5B). These distinct pH-dependent interactions suggest that TMs may undergo conformational rearrangements, leading to local structural changes that facilitate gate opening. Notably, the structural flexibility of TM2 and TM10 is consistent with their established role in the arginine transport cycle of a related transporter, AdiC arginine transporter, where their movement modulates substrate occlusion within the transporter[32].

**Figure 5.**
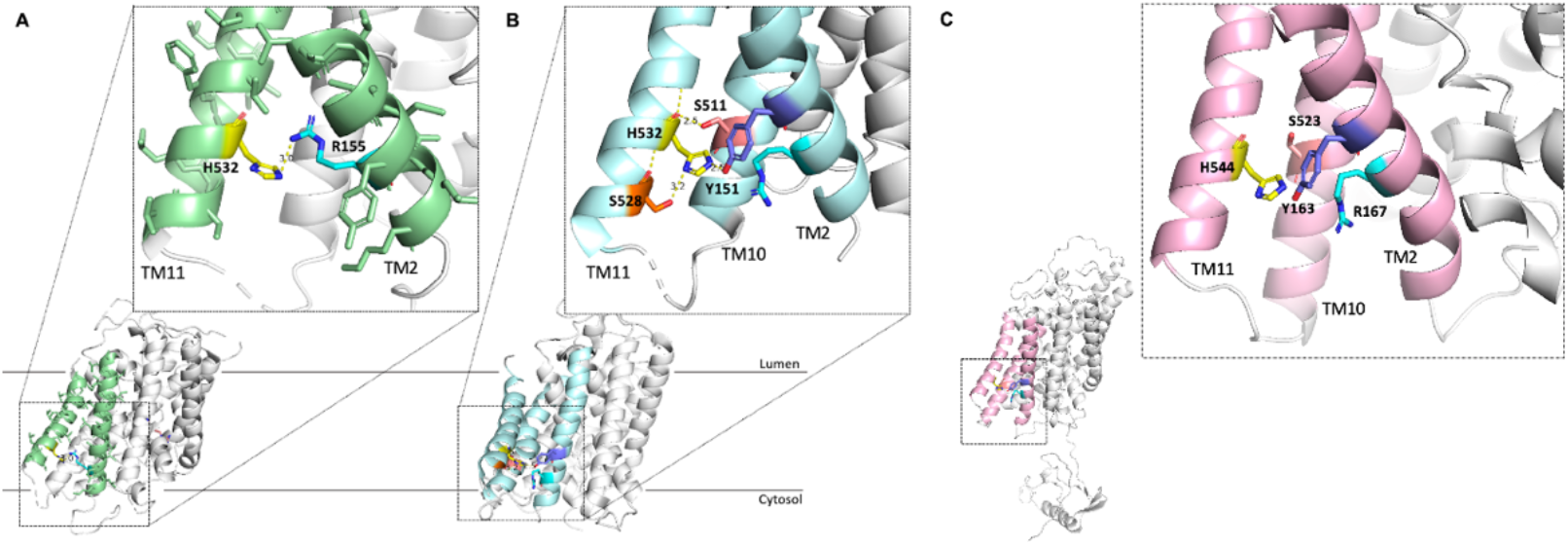
pH-sensing mechanism of His544 in hSLC38A9. The interactions between His532 and its surrounding residues in drSLC38A9 shift dynamically with the protonation and deprotonation of His532. **(A)** In the cytosol-open state structure of drSLC38A9 with bound substrate arginine (PDB: 6C08), deprotonated His532 interacts exclusively with Arg155 from TM2. His532 is shown in yellow, while Arg155 is shown in cyan. **(B)** In substrate arginine-free structure of drSLC38A9 (PDB: 7kgv), protonated His532 tightly interacts with various surround residues: Tyr151 (pink) from TM2, Ser511 (purple) from TM10, Ser528 (orange) from TM11. The Arg155 as shown in cyan was too far away from interacting with His532. **(C)**The AlphaFold model of hSLC38A9 reveals that His544 is surrounded by conserved residues that may undergo interaction shifts with His544 during its protonation and deprotonation. His544 is shown in yellow, Tyr163 in purple, Arg167 in cyan, and Ser523 in pink.

Given the high sequence conservation between hSLC38A9 and drSLC38A9, it is likely that His544 undergoes analogous protonation-dependent interaction shift, facilitating conformational transitions in hSLC38A9. Structural modeling using AlphaFold supports this postulate, showing that His544 in hSLC38A9 is surrounded by Arg167 and Tyr163 from TM2, and Ser523 from TM10 (Figure 5C). This suggests that under high-pH conditions, His544 may preferentially interact with Arg167, whereas under low-pH conditions, it may form interactions with Tyr163 and Ser523. Furthermore, the conservation of His544 and its associated interactions across species indicates that this histidine residue functions as a conserved pH sensor in SLC38A9 homologs.

## DISCUSSION

The lysosome is a membrane-bound organelle responsible for degrading biological macromolecules delivered through endocytic, phagocytic, and autophagic pathways[33,34]. Its lumen contains more than 60 hydrolases, most of which function optimally in an acidic environment[35,36]. To support their activity, the lysosomal pH is tightly maintained between 4.5-5.0[37,38], in contrast to the cytosolic pH of 7.0-7.5[39,40]. This steep gradient is established and sustained by the vacuolar ATPase (V-ATPase), which actively pumps protons into the lysosome[41,42]. Beyond enabling hydrolase function, the 100-1000-fold proton gradient also provides the driving force for many lysosomal exporters[42,43]. Indeed, numerous transporters localized to lysosomal and late endosomal membranes exhibit strong pH dependence, as the acidic lumen regulates their structural conformation, substrate affinity, and transport activity[20,44–46].

For example, cystinosin (SLC66A1), a lysosomal transporter that exports cystine into the cytosol, relies on a highly pH-dependent, proton-coupled mechanism. pH changes induce conformational shifts in cystinosin, suggesting that protonation of specific residues facilitates transport[47]. Notably, not all lysosomal transporters couple their activity directly to the proton gradient. PQLC2, a PQ-loop repeat-containing protein that mediates efflux of cationic amino acids, also requires the acidic environment for activity[48], but its transport is uncoupled from the proton gradient[4].

SLC38A9, a low-activity lysosomal arginine transporter, differs from classical high-capacity transporters in that its relatively weak transport efficiency enables it to serve as both a transporter and a nutrient sensor (“transceptor”) for mTORC1 activation[14]. This dual role necessitates tight regulation by the intracellular environment. To investigate how such regulation is achieved, we focused on the influence of pH on SLC38A9 activity and sought to identify the molecular determinant responsible for this pH-dependent modulation.

Here we demonstrate that SLC38A9 is activated by pH but relies on the sodium gradient to drive transport. Our study identified His544 as the critical residue responsible for pH sensing in SLC38A9. Using site-directed mutagenesis, we systematically mutated conserved histidine residues. Arginine uptake assays showed that three of the four conserved histidines (His339, His373, and His480) are not required for pH-dependent activity, as their mutants retained pH-sensitive transport (Supplemental Figure 2). In contrast, His544 is essential: the H544A mutant completely lost pH sensitivity. Similarly, a quadruple histidine mutant (4H) lacking all four conserved histidines showed no pH dependence, whereas reintroduction of His544 (3H mutant) restored pH-dependent arginine uptake (Supplemental Figure 2). Importantly, His544 primarily mediates pH regulation without substantially affecting transport itself, as both the H544A and 4H mutants retained baseline arginine transport activity. These findings indicate that His544 does not directly participate in substrate binding but instead functions as a pH-sensitive regulatory site. Thus, local ionic changes across the lysosomal membrane likely regulate arginine transport by inducing allosteric changes in SLC38A9.

We propose that His544 acts as an intrinsic pH sensor that allosterically modulates the transport activity of SLC38A9. At the neutral pH (7.2), the imidazole ring of His544 is unprotonated. In this condition, the aromatic ring stabilizes nearby protonated side chains through cation-π stacking, most prominently with the conserved Arg167 in the human protein (Arg155 in the zebrafish orthologue)[22]. This contact anchors transmembrane helices 2 and 10 in a conformation that favors substrate binding (Figure 5A).

When the lumen acidifies to pH 6.0, His544 becomes protonated and gains a positive charge (Figure 6). The former attractive interaction with Arg167 turns repulsive and the original clamp is released[22]. Protonated His544 can now form new cation-π or hydrogen bond contacts with neighboring aromatic or polar residues - Tyr163 and Ser523 in the human protein, Tyr151 and Ser511 in the zebrafish protein - which realigns transmembrane helices 2 and 10 into the inward open, substrate-free conformation observed at the low pH (Figure 5B). A similar mechanism was proposed for other amino acid transporters such as AdiC[49], SNAT2 and SNAT5[31].

**Figure 6.**
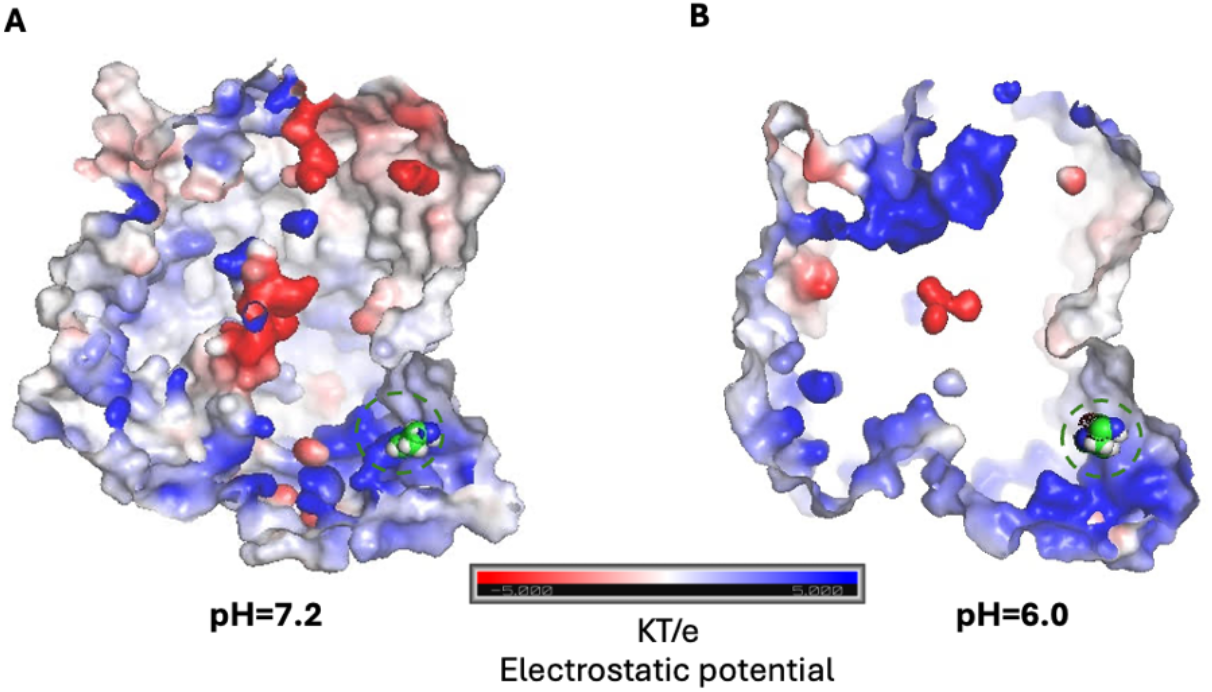
Comparison of drSLC38A9 electrostatic surface potential at high and low pH, revealing the pH-sensing role of His532. Electrostatics maps for drSLC38A9 structure in different pH. **(A)** Electrostatic surface of the drSLC38A9 under high pH; **(B)** Electrostatic surface of the drSLC38A9 under low pH. Locations of His532 are outlined by a dashed green circle.

Because these two transmembrane helices form the principal gates that occlude or release arginine during the transport cycle (as seen in AdiC[49]), this pH-dependent interaction modulation provides a direct molecular link between luminal acidity and transport kinetics. The conservation of His544 and its Arg, Tyr, and Ser partners across SLC38A9 homologues supports an evolutionarily preserved mechanism. Alternating protonation states of His544 therefore rewire its local interaction network and drive the structural transitions that underlie arginine uptake and signaling by SLC38A9. Since pH serves as a proxy for cellular metabolism, fluctuations in intracellular pH often reflect changes in metabolic state[50]. Indeed, altered intracellular pH is strongly associated with aging and numerous human diseases, including neurodegenerative disorders and cancer[51–53]. By sensing pH, SLC38A9 is positioned to integrate metabolic signals and adjust nutrient uptake in accordance with the cell’s physiological state.

### Experimental procedures

#### Generation of SLC38A9 variants

The sequence encoding hSLC38A9 (Genbank<u>Q8NBW4</u>) was cloned into the *p*Fastbac 1 vector (Invitrogen) with a sequence encoding a C-terminal 8X His tag and a thrombin cleavage site. Site-directed mutagenesis was carried out using a standard protocol involving two PCR steps. All SLC38A9 histidine mutants were confirmed by whole-plasmid sequencing. Verified plasmids were transformed into DH10Bac cells for preparation of bacmids. Recombinant baculovirus was generated and used for transfection following the protocol for the Bac-to-Bac Baculovirus Expression System (Invitrogen). Wild-type SLC38A9 and histidine mutants were overexpressed in *Spodoptera frugiperda* Sf-9 insect cells, which were harvested at 60 hours after P2 virus infection.

#### Protein Preparation

Harvested cell pellets were resuspended in lysis buffer containing 20mM Tris (pH 8.0) and 150mM NaCl and supplemented with cOmplete Protease inhibitor Cocktail (Roche), and 10mM EDTA, 10mM EGTA, 10mM DTT. Eighty homogenizing cycles were then carried out to break cells on ice, followed by ultracentrifugation at 130,000 x *g* for 40 mins. The pelleted membrane was then resuspended and washed with the high-high salt buffer containing 20mM Tris pH 8.0 and 1.6M NaCl followed by ultracentrifugation at 130,000 x *g* for 30 mins. The wash step was repeated twice. The pellets were then resuspended in the Lysis buffer, frozen in liquid nitrogen and stored at -80°C until further use.

To purify SLC38A9 protein and its variants, the membrane fraction was thawed and solubilized with 2% n-dodecyl-β-D-maltopyranoside (DDM, Anatrace), 0.2% Cholesteryl Hemisuccinate Tris salt (CHS, Anatrace) in 20 mM Tris pH 8.0, 160mM NaCl with 10% glycerol for 2 hours at 4°C. Following another ultra-centrifugation at 130,000 x *g* for 45 mins, the supernatant was loaded onto TALON Metal Affinity Resin (Clontech) and incubated at 4 °C overnight. The resin was washed by 25X column volumes of 20mM imidazole in a high-salt buffer containing 20mM Tris pH 8.0, 500 mM NaCl, 0.2% DDM. Then the protein was eluted with an elution buffer containing 300mM imidazole, 0.4% decyl-β-D-maltoside (DM), 0.02% DDM, 20mM Tris pH 8.0 and 160mM NaCl. The eluted proteins were applied to a Superdex-200 column (GE Healthcare) in a buffer containing 20mM Tris 8.0, 150mM NaCl, and 0.2% DM. The peak fraction was collected and concentrated for proteoliposome reconstitution and transport assays.

#### Preparation of Proteoliposomes

Chloroform-dissolved chicken egg phosphatidylcholine (egg-PC, Avanti Polar Lipids) was evaporated using dry nitrogen to yield a lipid film in a small glass vial and further dried under vacuum overnight. The lipids were hydrated in the inside buffer (20 mM Tris 8.0, 90 mM KCl, 10 mM NaCl) at 25 mg/mL by vortex for 3 minutes and then aged at room temperature for 1 hour. Liposomes were clarified by 5 rounds of freezing and thawing in liquid nitrogen and extruded through a 100 nm membrane with 21 passes (Millipore). The liposomes were pre-incubated with 1% n-octyl-b-D-glucoside (-OG) and 1 mM DDT for 1 hour at 4C before protein reconstitution. Purified wild-type SLC38A9 and variants were incorporated at a 1:100 (w/w) ratio into destabilized liposomes for 1 hour in the 4°C rotator. Glycerol-supplemented protein buffer was used in lieu of SLC38A9 protein in liposome-only control groups. The detergents were removed by incubation overnight with 200 mg per reaction Bio-Beads, and the proteoliposomes were further incubated with 40 mg per reaction fresh Bio-Beads for an additional hour. The proteoliposomes and liposome-only controls were collected using ultracentrifuge at 100,000Xg for 30 minutes at 4°C and then resuspended in an outside buffer (100 mM NaCl, different pH buffer: 20mM Mes 5.5, Mes 6.0, Mes 6.5, Tris 7.0, Tris 7.5 and Tris 8.2) to final lipid concentration of 40g/L.

#### Transport Assay

Transport reactions were initiated by adding 0.5M L-[^3^H]-arginine (American Radiolabeled Chemicals, Inc) to 50 L of proteoliposomes. Assays of Liposome-only controls were carried out in parallel to experimental groups as negative controls. All buffers were chilled, and assays were performed at a 30°C metal bath. For time-course uptake assay, at various time points, proteoliposomes were filtered, washed by 10mL of ice-cold wash buffer (outside buffer with 10 mM unlabeled L-arginine), and collected on 0.22 m nitrocellulose membranes (Millipore) which had been pre-wet by washing buffer. After washing, each membrane was dried by vacuum for exactly 1 minute and transferred into a glass vial with 10 mL scintillation fluid for counting. A time course profile indicates that the retained radio-ligands reached saturation after 10 mins. For the pH-dependent uptake assay, the equilibrium transport of arginine was measured at 10 min after adding 0.5M L-[^3^H]-arginine with proteoliposomes. The reaction was quickly placed into 0.22 m nitrocellulose membranes (Millipore) and washed with the 10mL ice-cold wash buffer (which had the same pH as the outside buffer with 10 mM unlabeled L-arginine). The membranes were collected into a glass vial for liquid scintillation counting. Non-specific adsorptions of L-[^3^H]-arginine by liposomes-only controls were subtracted from experimental measurements. All experimental and control groups were repeated two to three times.

## Data availability

All data are contained within the manuscript.

## Acknowledgments

We would like to thank Dr M. Gallenito (UCLA) for helpful discussions. This study was supported by the National Institutes of Health P41GM136508, RM1GM158451 and NCI F32CA278619. The Gonen laboratory is supported by funds from the Howard Hughes Medical Institute.

## Author contributions

All authors participated in the design of this project. X.M. performed protein expression, purification and proteoliposome reconstitution. X.M. performed functional experiments and data collection. X.M., A.S.H, and T.G. participated in data analysis and figure preparation. All authors wrote the manuscript.

## Funding and additional information

This study was supported by the National Institutes of Health P41GM136508, RM1GM158451 and NCI F32CA278619. The Gonen laboratory is supported by funds from the Howard Hughes Medical Institute.

## Supplementary Figures

**Supplemental Figure 1.**
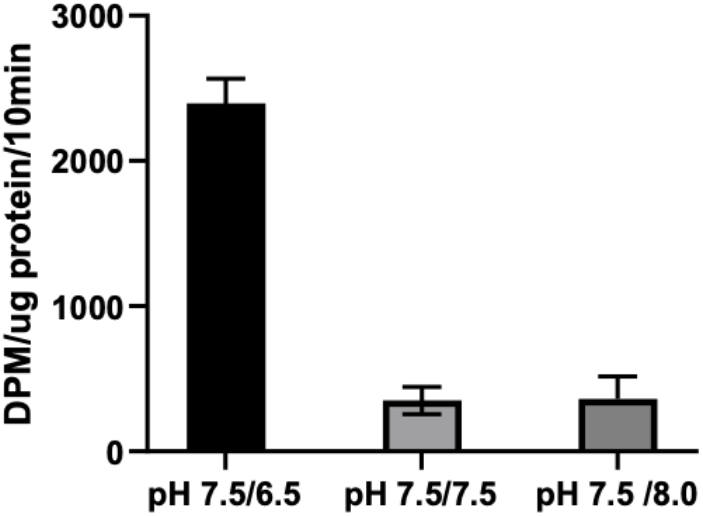
[^3^H]-arginine uptake activity of drSLC38A9 proteoliposomes in the presence of an inward vs outward H^+^ gradient vs symmetric pH. All the experiments had an inward Na^+^ gradient. Error bars, s.e.m. from three independent proteoliposome preparations; *n* = 3 biological replicates. Background uptake by liposome controls (without SLC38A9) were subtracted from these final arginine uptake graphs.

**Supplemental Figure 2.**
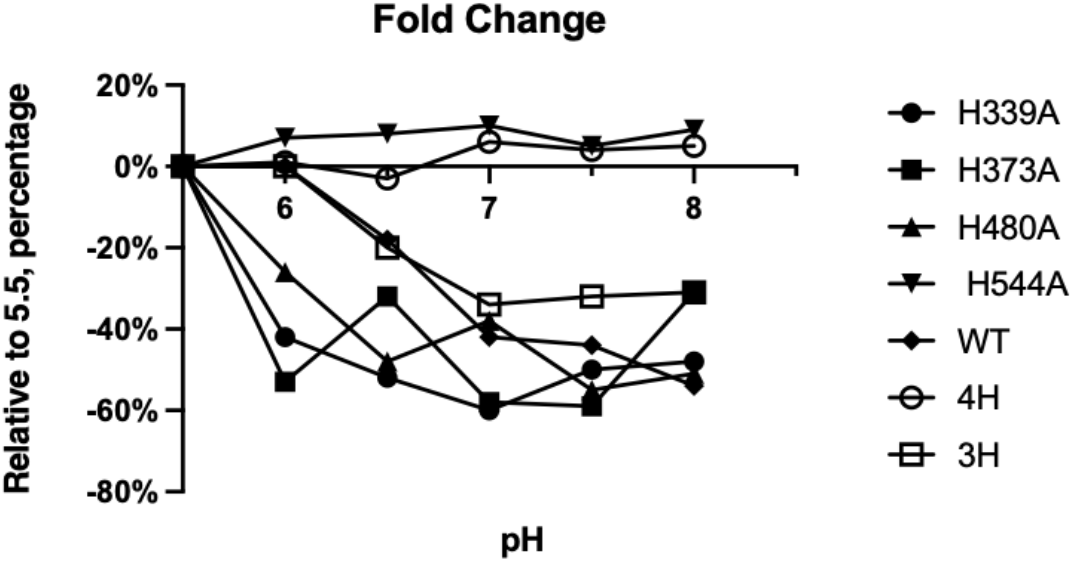
Analysis [^3^H]-arginine uptake assays of histidine mutants of SLC38A9. Uptake assays of single-site mutants of hSLC38A9 (H339A, H373A, H480A, and H544A) in proteoliposomes are compared to pH 5.5 and plotted as percentage changes in the y-axis. 4H mutant represents the mutant where all 4 histidines were mutated to alanine (H339A, H373A, H480A, H544A). 3H mutant represents the mutant where only 3 histidines were mutated to alanine (H339A, H373A, H480A). The results show the fold change of [^3^H]-arginine uptake at different pH compared to its at pH5.5, in each mutant and the wild-type SLC38A9.

