## Supplemental figures for "pH-dependent regulation in SLC38A9"

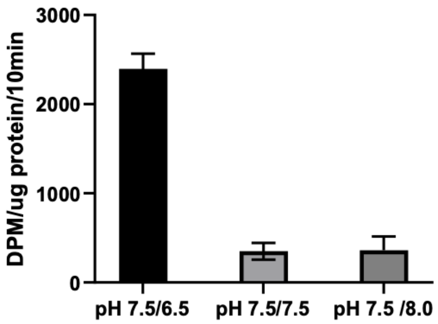


**Supplemental Figure 1. [$^3$H]-arginine uptake activity of drSLC38A9 proteoliposomes** in the presence of an inward vs outward $H^+$ gradient vs symmetric pH. All the experiments had an inward $Na^+$ gradient.
Error bars, s.e.m. from three independent proteoliposome preparations; *n* = 3 biological replicates. Background uptake by liposome controls (without SLC38A9) were subtracted from these final arginine uptake graphs.

Supplementary Figures:

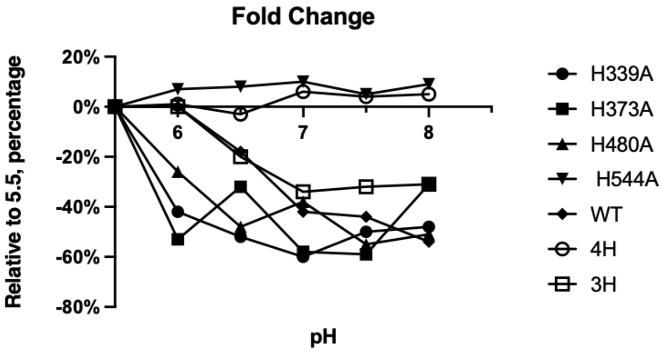


**Supplemental Figure 2. Analysis [$^3$H]-arginine uptake assays of histidine mutants of SLC38A9**. Uptake assays of single-site mutants of hSLC38A9 (H339A, H373A, H480A, and H544A) in proteoliposomes are compared to pH 5.5 and plotted as percentage changes in the y-axis. 4H mutant represents the mutant where all 4 histidines were mutated to alanine (H339A, H373A, H480A, H544A). 3H mutant represents the mutant where only 3 histidines were mutated to alanine (H339A, H373A, H480A). The results show the fold change of [$^3$H]-arginine uptake at different pH compared to its at pH5.5, in each mutant and the wild-type SLC38A9.
